# CRISPR-Cas13d screens identify *KILR*, a breast cancer risk-associated lncRNA that regulates DNA replication and repair

**DOI:** 10.1101/2023.11.16.567471

**Authors:** Lu Wang, Mainá Bitar, Xue Lu, Sebastien Jacquelin, Sneha Nair, Haran Sivakumaran, Kristine M. Hillman, Susanne Kaufmann, Rebekah Ziegman, Francesco Casciello, Harsha Gowda, Joseph Rosenbluh, Stacey L. Edwards, Juliet D. French

## Abstract

Long noncoding RNAs (lncRNAs) have surpassed the number of protein-coding genes, yet the majority have no known function. We previously discovered >800 lncRNAs at regions identified by breast cancer genome-wide association studies (GWAS). Here, we performed a pooled CRISPR-Cas13d RNA knockdown screen to identify which of these lncRNAs altered cell proliferation. We found that *KILR,* a lncRNA that functions as a tumor suppressor, safeguards breast cells against uncontrolled proliferation. The half-life of *KILR* is significantly reduced by the risk haplotype, revealing an alternative mechanism by which variants alter cancer risk. We showed that *KILR* sequesters RPA1, a subunit of the RPA complex, required for DNA replication and repair. Reduced *KILR* expression promotes cell proliferation by increasing the available pool of RPA1 and the speed of DNA replication. Our findings confirm lncRNAs as mediators of breast cancer risk, emphasize the need to annotate noncoding transcripts in relevant cell types when investigating GWAS variants and provide a scalable platform for mapping phenotypes associated with lncRNAs.

## INTRODUCTION

Genome-wide association studies (GWAS) have identified thousands of genetic variants associated with normal and disease traits. Most variants are located in noncoding regions of the genome and do not directly affect protein-coding sequences. A well-established mechanism by which GWAS variants modulate disease risk is through the alteration of DNA enhancers, causing changes in the expression of nearby target genes^1,2^. In addition to housing DNA regulatory elements, the human noncoding genome is pervasively transcribed and at least a subset of the resulting molecules are functional at the transcript level. The majority of the transcripts are long noncoding RNAs (lncRNAs), defined as RNA transcripts longer than 200 nucleotides that do not code for proteins.

Despite lacking the ability to code for proteins, lncRNAs perform a diverse array of cellular functions^3^. In the nucleus, they can alter transcription by guiding epigenetic modifications and/or transcription factors, regulate splicing by binding splicing factors, act as scaffolds for protein complexes and promote the formation of nuclear bodies and domains^4^. In the cytoplasm, lncRNAs have been shown to regulate RNA stability and translation, interact with proteins to affect their localization, stability, and post-translational modifications and influence cellular export and signaling pathways, among other described functions^5^. LncRNAs display exquisite cell-type specific expression and are important in defining specific cell subpopulations and cell states^6^. The aberrant expression of lncRNAs has been reported in various disease phenotypes, including cancer, and many have been directly implicated in disease development^3^. However, the impact of disease risk-associated variants on lncRNA expression and function is less evident and requires further study.

We recently discovered thousands of lncRNAs transcribed from breast cancer GWAS loci and nearby regions^7^ (<1.5 Mb). An enrichment of GWAS variants was observed in lncRNA exons but not in their introns or promoter regions, suggesting that lncRNA transcripts are important mediators of breast cancer risk. We identified 844 lncRNAs as potential GWAS target genes based on the presence of breast cancer risk variants in their exons, promoters or distal DNA regulatory elements^7^. Expression quantitative trait loci (eQTL) analyses identified lncRNAs whose expression are associated with risk variants in breast tumors, providing additional evidence for their involvement in breast cancer development^7^. From our findings, we expected that some of the identified lncRNAs would influence breast cell proliferation.

High-throughput pooled loss-of-function screens are a powerful strategy for identifying genes implicated in different phenotypes. CRISPR-Cas9 cutting (CRISPRko), targets DNA regions in the genome and is the most used strategy for protein-coding gene knockouts. CRISPRko will often be ineffective for lncRNAs as the cutting may not alter lncRNA stability or function. Several CRISPR-dCas9-based activation (CRISPRa) and inhibition (CRISPRi) screens have successfully been used to overexpress and knockdown mRNA and lncRNAs^8–11^. However, given that lncRNA transcription is often initiated from enhancer elements encoded in DNA, it is not clear whether the observed CRISPRi/a effect is DNA or RNA mediated. To overcome these hurdles, we performed CRISPR-Cas13d RNA knockdown screens to identify the breast cancer-associated lncRNAs whose knockdown affects proliferation of normal breast and breast cancer cells.

## RESULTS

### Breast cancer-associated lncRNAs can alter cell proliferation

To identify breast cancer risk-associated lncRNAs that regulate cell proliferation, CRISPR-Cas13d-based knockdown screens were performed in a normal mammary epithelial cell line^12^ (K5+/K19+) and two breast cancer cell lines (estrogen receptor, ER-positive MCF7 cells and ER-negative MDAMB231 cells; **Fig. 1a**). A pooled CRISPR-Cas13 guide RNA (crRNA) library targeting 1864 lncRNAs and protein-coding genes was designed, aiming for ten cRNAs predicted to be of high-quality per gene. However, for 5% of the genes targeted, only two to nine guides met our quality control criteria (see Methods). The gene set included, i) the 844 breast cancer-associated lncRNAs we previously linked with GWAS risk variants^7^; ii) 59 annotated lncRNAs; and iii) 935 protein-coding genes, including (with overlap) 129 estrogen-regulated genes^13^, 246 breast cancer driver genes^14^ and 612 essential genes^14^. In addition, crRNAs targeting ten protein-coding genes and ten lncRNAs from chromosome Y were included as negative controls. In total, the synthesized library contained 18,248 crRNAs (**Supplementary Table 1**).

**Figure 1.**
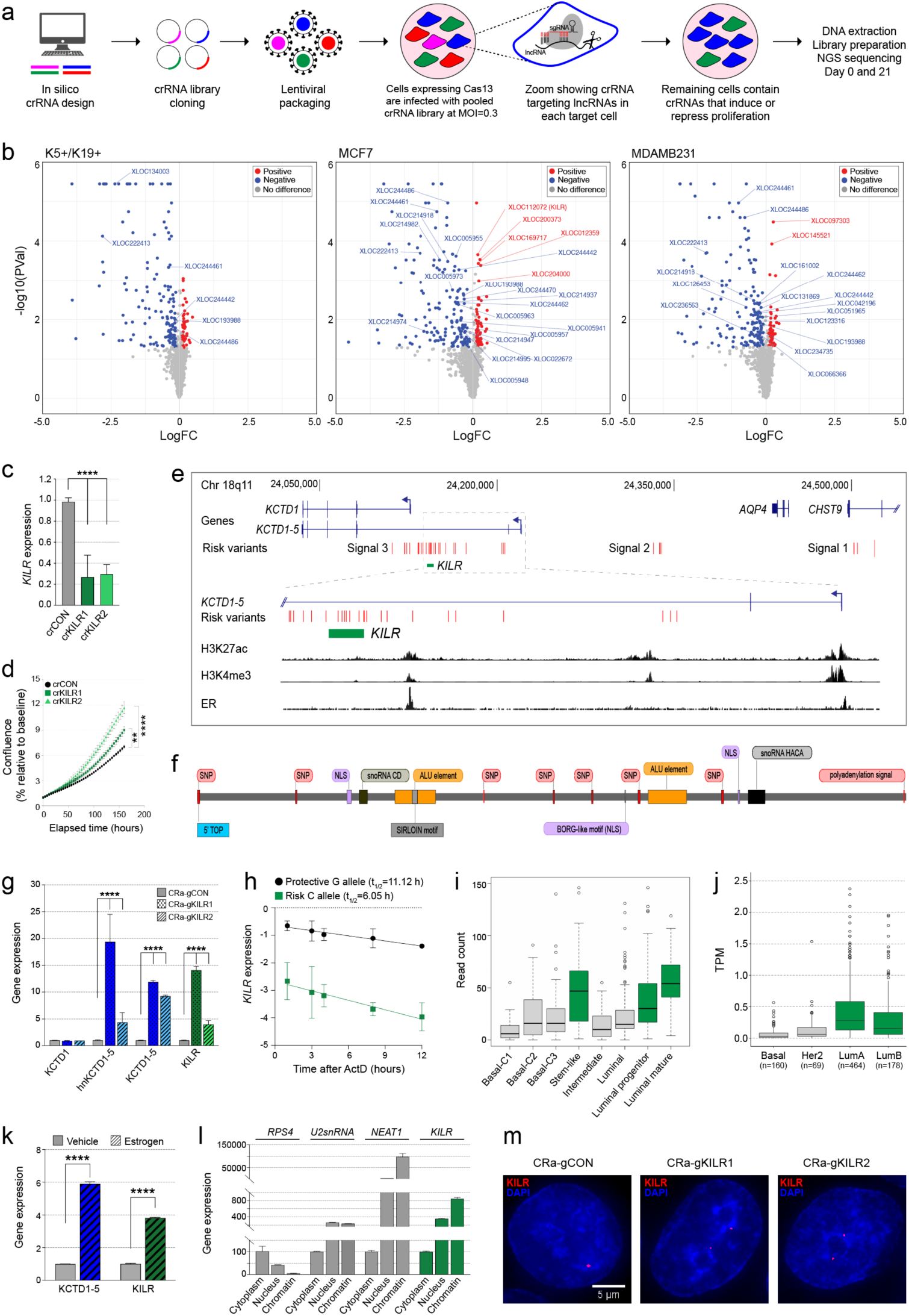
CRISPR-Cas13d screens identify lncRNAs that modulate breast cell proliferation. (**a**) Schematic for CRISPR-Cas13d screens. (**b**) Scatterplots from CRISPR-Cas13d screen data showing differentially represented crRNAs (red/blue dots; log2[fold-change] > 0.1 and p value < 0.05) targeting candidate genes and lncRNAs. Labels are unannotated breast cancer-associated lncRNAs with FDR ≤ 0.3. (**c**) qPCR for *KILR* expression and (**d)** cell confluence measured over time using Incucyte in MCF7 cells after Cas13d-*KILR* knockdown with two independent crRNAs (crKILR1-2). The crCON contains a non-targeting control. Error bars, SEM (n = 3). p values were determined by one-way ANOVA and Dunnett’s multiple comparisons test (**p < 0.01, ****p < 0.0001). (**e**) WashU genome browser (hg19) showing GENCODE annotated genes (blue) and *KILR* (green). The breast cancer risk variants are shown as red vertical lines (Signals 1-3). The H3K27ac, H3K4me3 and ER (estrogen receptor) binding tracks from MCF7 cells are shown as black histograms. (**f**) A linear schema of *KILR*. SNP (single nucleotide polymorphism); 5’ TOP (terminal oligopyrimidine tract); NLS (nuclear localization signal). (**g**) qPCR for *KCTD1, KCTD1-5* and *KILR* expression in T47D cells after CRISPRa activation of the *KCTD1-5* promoter to overexpress *KILR* with two independent gRNAs (CRa-gKILR1-2). The CRa-gCON contains a non-targeting control. Error bars, SEM (n = 3). p values were determined by one-way ANOVA and Dunnett’s multiple comparisons test (****p < 0.0001). (**h**) *KILR* RNA stability assay in MDAMB361 cells after treatment with actinomycin D (ActD), then qPCR for *KILR* RNA relative to *CDKN2A* mRNA levels. *KILR* mRNA half-life (t_1/2_) was calculated by linear regression analysis. Error bars, SEM (n = 3). (**i**) Boxplot of *KILR* read counts in normal breast tissue from scRNA-seq data clustered based on NB-lncRNA expression^6^. (**j**) Boxplot of *KILR* TPM (transcript per million) in breast tumour samples from TCGA RNA-seq data stratified by tumour subtype. (**k**) qPCR for *KCTD1-5* and *KILR* expression in estrogen-stimulated T47D cells. Error bars, SEM (n = 3). p values were determined by two-way ANOVA and Dunnett’s multiple comparisons test (****p < 0.0001). (**l**) qPCR after nuclear/cytoplasmic/chromatin fractionation of T47D cells detecting the distribution of the indicated transcripts. Error bars, SD (n = 2). (**m**) Representative confocal microscopy images of *KILR* in MCF7 cells after CRISPRa (CRa-gKILR1-2) stained with Stellaris *KILR* RNA FISH probes (red). The CRa-gCON contains a non-targeting control. Nuclei were stained with DAPI (blue). Scale bar, 5 μm.

Breast cells expressing CRISPR-Cas13d were infected with lentivirus expressing the crRNA library (multiplicity of infection, MOI=0.3) and antibiotic selected for four days (**Fig. 1a**). Three biological replicates were generated for every cell line, each with independent crRNA library transductions. After plating, cells were cultured for 21 days, collected for DNA extraction and crRNA abundance was quantified by next generation sequencing. Attesting to the quality of the screen, we observed a high correlation (>0.9) between replicates and a low Gini index (<0.1), indicating even crRNA read counts, successful transduction and absence of over-selection (**Supplementary** Figs. 1a-d). Moreover, none of the negative controls had significant effects on cell proliferation. The magnitude of effect (Log2[fold-change]) and statistical significance of each gene in the screen were calculated using MAGeCK^15^ based on the estimated abundance of all targeting crRNAs at the start and end of the experiment. To further reduce the effect of off targets and increase the confidence of the screens, we filtered out crRNAs with full or partial (up to two mismatches and down to 90% coverage) complementarity to additional genome or transcriptome regions from the *in silico* crRNA library (**Supplementary Table 1**). After running MAGeCK based on the filtered crRNA library, lncRNAs with FDR <=0.3 were considered as high confidence and are referred to as positive (knockdown increased cell proliferation) or negative (knockdown decreased cell proliferation) hits (**Fig. 1b**).

As expected, knockdown of essential genes (e.g. *EIF3B*, *RPL35*) and oncogenes (e.g., *AKT1*, *CDH1*, *EGFR*, *MYC*) had a negative effect on cell proliferation, whereas knockdown of tumor-suppressor genes (e.g. *TP53, EP300, CASP8)* had a positive effect (**Fig. 1b** and **Supplementary Table 2**). Annotated lncRNAs already implicated in breast cancer (*DSCAM-AS1*, *NEAT1*, *SNHG3*, *ZFAS1*) also had a significant (FDR ≤ 0.3) effect, indicating the screen was able to identify functional lncRNAs. Additionally, knockdown of 39 high-confidence breast cancer-associated lncRNAs either suppressed or promoted proliferation, including five lncRNAs common to all three cell lines (**Supplementary** Fig. 1e). The effect of the best-performing crRNAs targeting five breast cancer-associated lncRNAs characterized as significant hits were individually validated in MCF7 cells, confirming the quality of the screen (**Supplementary** Figs. 1f,g and **Supplementary Table 2**).

### Breast cancer eQTL lncRNAs alter cell proliferation

Seven breast cancer-associated lncRNAs targeted in the screen were previously identified as eQTLs for breast cancer risk variants^7^. Four of these lncRNAs were hits in at least one of our CRISPR-Cas13d-based screens. Two of these eQTL lncRNAs, *XLOC209276* (a positive hit in MCF7 cells) and *XLOC022678* (a negative hit in MDAMB231 and K5+/K9+ cells), were initial hits that did not maintain significance (FDR<=0.3) after crRNA library filtering (**Supplementary Table 2**). Targeted knockdown of another two eQTL lncRNAs (*XLOC112072* and *XLOC169717*) resulted in significantly higher cell proliferation (positive hits) in MCF7 cells. Given its predicted function in cell proliferation, the absence of off-target effects, and genetic evidence of its role in breast cancer, we decided to prioritise lncRNA *XLOC112072* (hereafter named *KILR; <u>K</u>CTD1 Intronic <u>L</u>nc<u>R</u>NA*) for functional studies.

### *KILR* is a sense intronic lncRNA transcribed from the *KCTD1-5* promoter

The two most significant crRNAs targeting *KILR* were individually validated, confirming that both crRNAs can knock down *KILR* (**Fig. 1c**) and increased the proliferation of MCF7 cells (**Fig. 1d**). We used 5’ and 3’ random amplification of cDNA ends (RACE) to map the transcriptional ends of *KILR*, revealing a 6.7 kilobase (kb), single exon, polyadenylated transcript (**Fig. 1e, Supplementary File 1** and **Supplementary** Figs. 2a,b). *KILR* is located on chromosome 18q11 within the first intron of *KCTD1-*5, a transcript variant of *KCTD1* generated from an alternative transcription start site (TSS) located upstream of the canonical *KCTD1* promoter (**Fig. 1e**). *KILR* contains seven breast cancer risk variants, a 20 nucleotide 5’ poly(U) tract (commonly referred to as a terminal oligopyrimidine tract; 5’TOP), two predicted H/ACA box snoRNAs, a predicted C/D box snoRNA, a BORG-like motif, a SINE-derived nuclear localization (SIRLOIN) motif, multiple nuclear localization signals (NLS), and a ‘tailsout’ inverted repeat Alu element (IRAlu), known to promote lncRNA nuclear localization^16^ (**Fig. 1f**). Structural predictions of *KILR* confirm the formation of a long double strand, resulting from the pairing of the IRAlu elements, which confers upon *KILR* the characteristic structure of IRAlu-containing lncRNAs^17^ (**Supplementary** Fig. 2c). The 5’ end of *KILR* transcript does not coincide with histone modifications typically associated with promoter activity (i.e. H3K27Ac, H3K4Me3; **Fig. 1e**), thus we hypothesized that *KILR* may be generated from the upstream *KCTD1-5* promoter. To confirm this, we activated the *KCTD1-5* promoter by directing dCas9 fused to transcriptional activators (CRISPRa) to the TSS of *KCTD1-5* using two independent CRISPR-dCas9 guide RNAs (gRNAs) and showed increased expression of both *KCTD1-5* and *KILR*, but not *KCTD1* (**Fig. 1g**).

### Breast cancer risk variants at 18q11 reduce the half-life of *KILR*

Fine mapping of breast cancer GWAS data has identified three independent signals at 18q11 (**Fig. 1e**). We previously showed that genetic variants in Signal 3 are associated with reduced *KILR* expression in breast tumors and that the eQTL signal colocalizes with the risk signal^7^. Since seven highly correlated variants from Signal 3 fall within the *KILR* transcript, we hypothesized that the risk alleles may affect *KILR* expression by altering its RNA stability. To test this, we measured the allele-specific half-life (t_1/2_) of *KILR* following treatment with actinomycin D in MDAMB361 cells, a breast cancer cell line that is heterozygous for the breast cancer risk alleles. The t_1/2_ of *KILR* carrying the risk alleles is almost half that of *KILR* in the presence of the protective alleles (11 h compared with ∼6h; **Fig. 1h**). No allele-specific difference in RNA stability was observed for *KCTD1-5* hnRNA (**Supplementary** Fig. 2d). These results are consistent with our eQTL study and suggest that inclusion of the risk alleles reduces the expression of *KILR* by altering its transcript stability. Using capture Hi-C data that we generated for an independent study^18^, we showed that the regions containing the other two GWAS signals at 18q11 (Signal 1 and Signal 2) physically interact with the *KCTD1-5*/*KILR* promoter region in breast cell lines through chromatin looping (**Supplementary** Fig. 2e). These results suggest that additional nearby GWAS signals at 18q11 may also affect *KILR* (and *KCTD1-5*) expression. Overall, our observations are consistent with *KILR* being at least one of the target genes of the breast cancer GWAS signals at 18q11.

### *KILR* is a widely-expressed, estrogen-responsive nuclear-retained lncRNA

We confirmed that *KILR* is expressed in normal breast tissue (**Fig. 1i**), breast tumors (**Fig. 1j**) and normal breast and breast cancer cell lines (**Supplementary** Fig. 2f). In normal breast tissue, *KILR* expression is higher in cells of the luminal subtype (**Fig. 1i**), specifically in stem-like cells and in subpopulations that were significantly correlated with the PAM50 breast tumor subtypes^6^. In TCGA breast tumors, *KILR* is expressed at higher levels in the ER-positive breast cancer subtypes (luminal A and B; **Fig. 1j**). Both *KCTD1-5* and *KILR* are also induced by estrogen treatment (**Fig. 1k**), consistent with the presence of estrogen receptor binding at the *KCTD1-5* promoter (**Fig. 1e**). Given the presence of NLS and SIRLOIN motifs and the IRAlu element in its sequence (**Fig. 1f**), all of which promote nuclear retention^19^, we predicted *KILR* localization to be restricted to the nucleus. To confirm this, we performed subcellular fractionation of breast cancer cells, which showed that *KILR* is predominantly nuclear and enriched in the chromatin fraction (**Fig. 1l**). Furthermore, RNA-FISH following CRISPRa-induced *KILR*/*KCTD1-5* expression in MCF7 cells revealed that *KILR* formed distinct nuclear puncta (**Fig. 1m** and **Supplementary** Fig. 2g).

### *KILR* overexpression induces breast cancer cell apoptosis

Since the knockdown of *KILR* promotes cell proliferation, we sought to determine whether its overexpression would have the opposite effect. We induced *KILR* and *KCTD1-5* expression in one normal (MCF10A) and two cancer (MCF7s and Hs578T) breast cell lines, by targeting CRISPRa to their shared promoter (**Supplementary** Fig. 2a) and observed significantly suppressed cell proliferation in all three cell lines (**Fig. 2a**). In fact, we observed cell death a few days after induction, suggesting that increased expression of *KILR/KCTD1-5* may lead to apoptosis. Annexin V staining confirmed that CRISPRa-mediated induction of *KILR/KCTD1-5* promoted apoptosis in the two breast cancer cell lines, but not in the normal breast cell line (**Figs. 2b,c**). CRISPRa induction targets both *KILR* and *KCTD1-5*, thereby posing a challenge to pinpoint which of the two genes is promoting apoptosis. To address this, we used a Tet-On inducible lentiviral overexpression system to overexpress *KCTD1-5* or *KILR* (**Supplementary** Fig. 3b), and showed that individual overexpression of *KILR* but not *KCTD1-5* promoted apoptosis, again only in the breast cancer cell lines (**Figs. 2d,e**). *KILR*-induced apoptosis in breast tumor cells but not normal breast cells was also observed in additional cell lines (**Supplementary** Figs. 3c-e).

**Figure 2.**
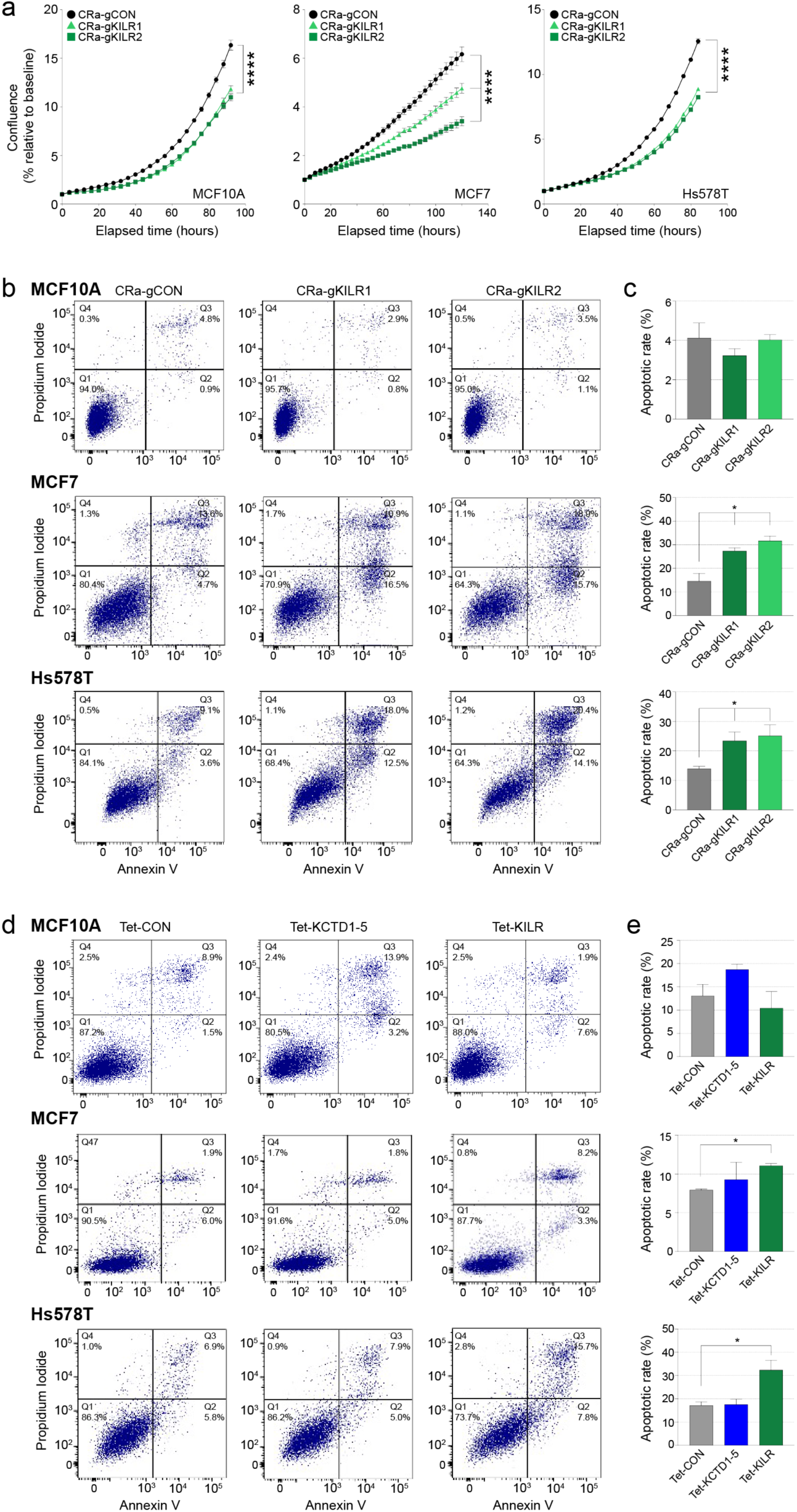
*KILR* overexpression inhibits breast cell proliferation and induces apoptosis. (**a**) Cell confluence measured over time using Incucyte in breast cells after CRISPRa activation of the *KCTD1-5* promoter to overexpress *KILR* with two independent gRNAs (CRa-gKILR1-2). The CRa-gCON contains a non-targeting control. Error bars, SEM (n = 3). p values were determined by one-way ANOVA and Dunnett’s multiple comparisons test (****p < 0.0001). (**b**) Representative apoptosis analysis of breast cells after CRISPRa (CRa-gKILR1-2) by double staining with annexin V and PI. The CRa-gCON contains a non-targeting control. The quadrants (Q) were defined as Q1 = live (Annexin V– and PI-negative), Q2 = early stage of apoptosis (Annexin V-positive/PI-negative), Q3 = late stage of apoptosis (Annexin V– and PI-positive) and Q4 = necrosis (Annexin V-negative/PI-positive). (**c**) The percentage of cells in early and late stage apoptosis in each group (Q2 + Q3). Error bars, SEM (n = 3). p values were determined by one-way ANOVA and Dunnett’s multiple comparisons test (*p < 0.05). (**d**) Representative apoptosis analysis of breast cells after doxycycline induction of ectopic *KCTD1-5* or *KILR* expression by double staining with annexin V and PI. The Tet-CON represents an empty vector control. The quadrants were defined in (b). (**e**) The percentage of cells in early and late stage apoptosis in each group (Q2 + Q3). Error bars, SEM (n = 3). p values were determined by one-way ANOVA and Dunnett’s multiple comparisons test (*p < 0.05).

### *KILR* inhibits DNA replication by sequestering the RPA1 protein

To identify proteins that physically interact with *KILR*, we performed RNA pull-down in MCF7 cells using *in vitro* transcribed *KILR* followed by mass spectrometry. *KILR*-binding proteins identified by the presence of at least five peptides were sorted according to their enrichment over a *LacZ* control experiment (**Supplementary Table 3**). One of the most highly enriched proteins was RPA1, the main single stranded DNA (ssDNA) binding protein in humans and member of the heterotrimeric RPA complex essential for multiple processes in DNA metabolism, including DNA replication and damage repair^20^. Using RNA FISH combined with immunofluorescence (IF), we confirmed *KILR* and RPA1 colocalization and showed that overexpression of *KILR* sequesters RPA1 into nuclear puncta (**Fig. 3a** and **Supplementary** Figs. 4a,b), suggesting *KILR* may abrogate RPA function by reducing the levels of available RPA1. Of note, the *BORG* lncRNA has been shown to physically interact with RPA1 in breast cancer^21^, suggesting *KILR* may interact with RPA1 via the BORG-like motifs in its sequence. RPA binds ssDNA at DNA replication forks and enhances the assembly and recruitment of DNA polymerases, thus facilitating DNA replication^22^. We hypothesized that the increased proliferation of MCF7 cells following Cas13d-mediated knockdown of *KILR* could be a result of an increased pool of RPA1 becoming available for DNA replication. In support of this, we performed a DNA fiber assay at single-molecule resolution and showed that knockdown of *KILR* using two independent crRNAs promotes proliferation by increasing the speed of DNA replication (**Fig. 3b**).

**Figure 3.**
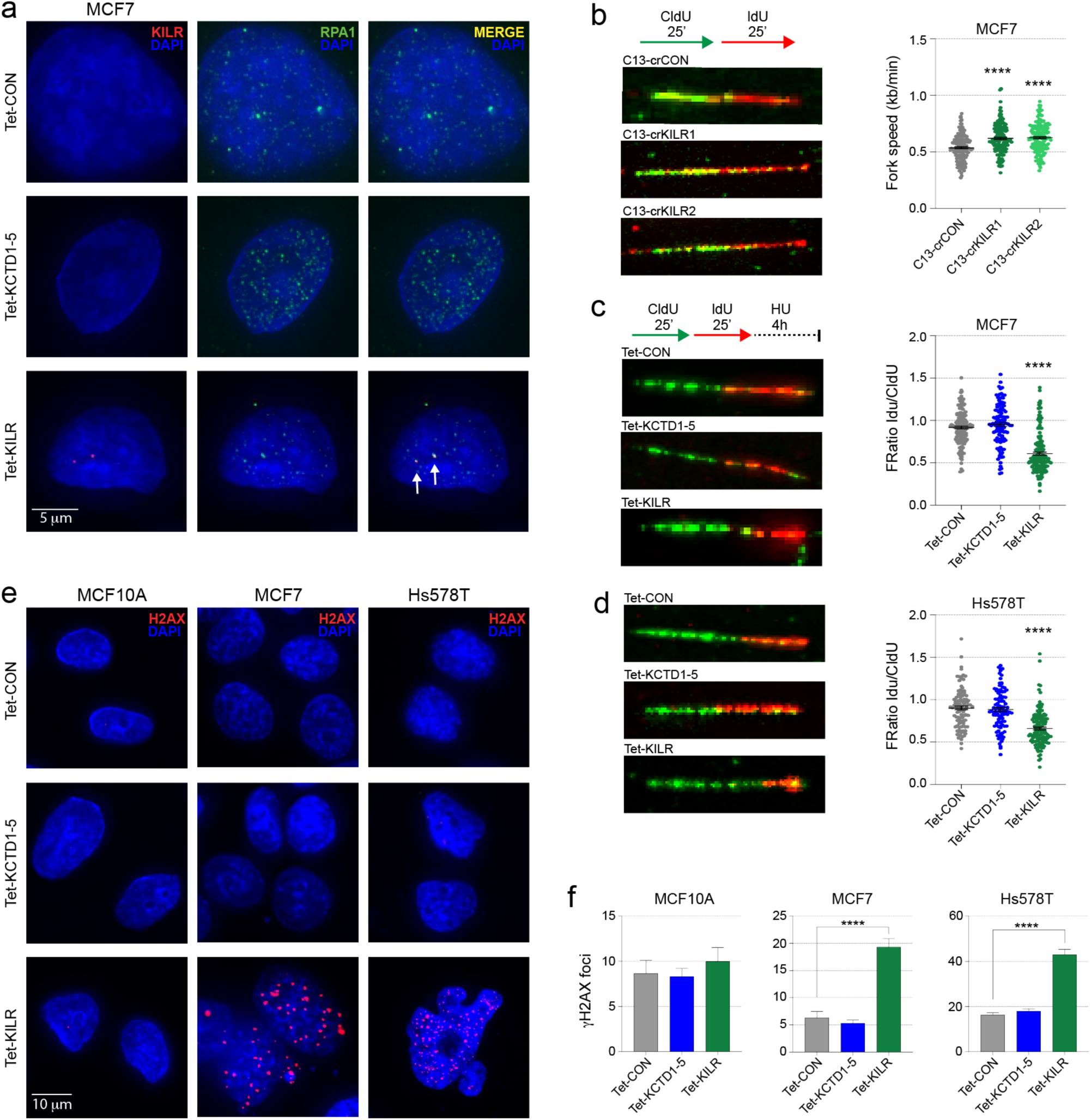
*KILR* inhibits DNA replication by sequestering the RPA1 protein. (**a**) Representative confocal microscopy images of *KILR* and RPA1 in MCF7 cells after doxycycline induction of ectopic *KCTD1-5* or *KILR* expression stained with Stellaris *KILR* RNA FISH probes (red) and immunostained with anti-RPA1 (green) (n = 3). The Tet-CON represents an empty vector control. Nuclei were stained with DAPI (blue). White arrows highlight *KILR*/RPA1 co-localization. Scale bar, 5 μm. (**b**) Left panels: Representative images of DNA fibers in MCF7 cells after Cas13d-*KILR* knockdown with two independent crRNAs (crKILR1-2) then labelling with CldU and IdU. Right panel: Replication fork speed was calculated by length of track/time of CIdU pulse. Data are presented from two independent fiber assays. Error bars, SEM (n = 154). p values were determined by one-way ANOVA and Dunnett’s multiple comparisons test (****p < 0.0001). (**c, d**) Left panels: Representative images of DNA fibers in MCF7 (**c**) and Hs578T (**d**) cells after doxycycline induction of *KCTD1-5* or *KILR*, labelling with CldU and IdU then treatment with 4 mM HU for 4 h. Right panels: Ratio of IdU/CldU. Data are presented from two independent fiber assays. Error bars, SEM (n = 150). p values were determined by one-way ANOVA and Dunnett’s multiple comparisons test (****p < 0.0001). (**e**) Representative confocal microscopy images of breast cells after doxycycline induction of ectopic *KCTD1-5* or *KILR* expression immunostained with anti-ɣH2AX (red). The Tet-CON represents an empty vector control. Nuclei were stained with DAPI (blue). Scale bar, 10 μm. (**f**) Quantification of ɣH2AX foci in three breast cell lines. A cell with > 10 distinct ɣH2AX foci in the nucleus was considered as positive. Error bars, SEM (n = 3). p values were determined by one-way ANOVA and Dunnett’s multiple comparisons test (****p < 0.0001).

More recently, it was shown that the RPA complex can stabilize the DNA replication fork through recruitment of the SWI/SNF family member, HARP^23,24^. Using a modified version of the DNA fiber assay, we showed that overexpression of *KILR* but not *KCTD1-5*, causes degradation of nascent DNA in MCF7 and Hs578T cells under hydroxyurea-induced replication stress (**Figs. 3c,d**), indicating fork protection defects. We also showed that MCF7 and Hs578T cells in which *KILR* has been overexpressed had increased DNA damage, marked by H2AX phosphorylation (into γH2AX), in the absence of DNA insult (**Figs. 3e,f**). The same phenotype was not observed in MCF10A cells or following *KCTD1-5* overexpression in all three cell lines (**Figs. 3e,f**). Taken together, these results suggest that by reducing the pool of available RPA1, overexpression of *KILR* in breast cancer cells mimics the phenotype of RPA1 knockdown.

### *KILR* inhibits homologous recombination-based repair

In addition to DNA replication, RPA also plays an important role in homologous recombination repair (HRR) of DNA double strand breaks (DSBs)^25^. A key step in the initiation of HRR is the generation of ssDNA through end resection. This ssDNA stretch is rapidly coated by RPA, which is subsequently replaced by RAD51^26^. In response to DNA damage caused by ionizing radiation (IR), overexpression of *KILR*, but not *KCTD1-5* inhibited RPA and RAD51 recruitment to DSBs in breast cancer cells (**Figs. 4a,b** and **Supplementary** Figs. 4c,d). Using RNA FISH and IF we showed that this effect is mediated by the sequestration of RPA1 to nuclear puncta (**Fig. 4c** and **Supplementary** Fig. 4e). Notably, a fraction of RPA1 is colocalized with *KILR* in nuclear foci in normal breast cells only (**Fig. 4d**), with the remaining being available for DNA damage repair. As the overexpression of *KILR* inhibited RPA and RAD51 mobilization to DSBs, we hypothesized that *KILR* knockdown would have the opposite effect, resulting in a more efficient DNA damage repair. Consistent with this, breast cancer cells in which *KILR* was knocked down using Cas13d resolved ∼50% of the IR-induced DSBs within the first six hours after irradiation (**Fig. 4e**). The same was not observed in the Cas13d-control cells where the number of DSBs remained constant. This indicates that HRR is more efficient in the absence of *KILR*, likely due to an increase in the pool of available RPA1.

**Figure 4.**
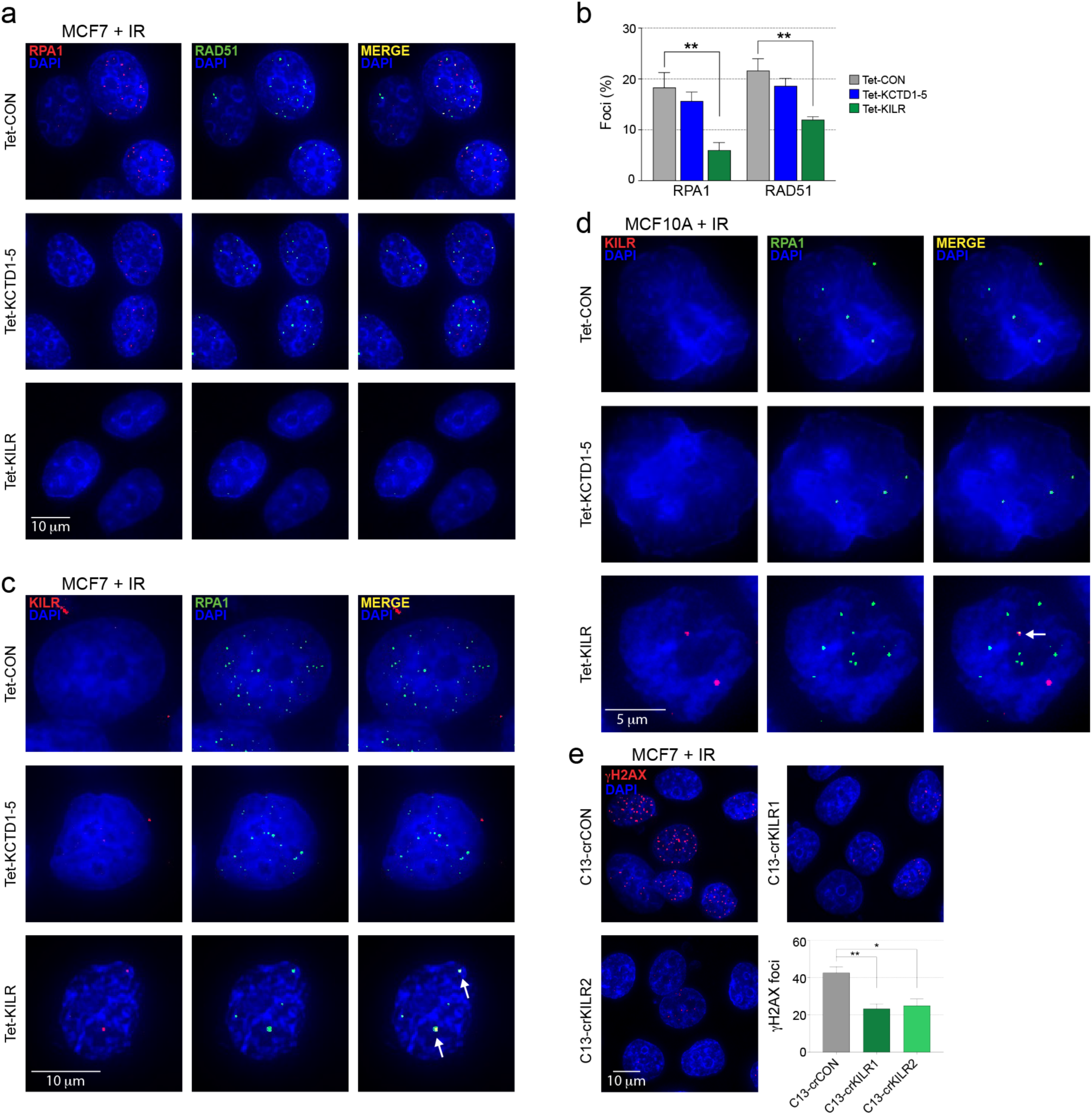
*KILR* overexpression inhibits HR-based repair. (**a**) Representative confocal microscopy images of RPA1 and RAD51 in MCF7 cells after doxycycline induction of ectopic *KCTD1-5* or *KILR* expression and exposure to 6-Gy IR (n = 3). 6 h post-IR, cells were immunostained with anti-RPA1 (red) and anti-RAD51 (green). The Tet-CON represents an empty vector control. Nuclei were stained with DAPI (blue). Scale bar, 10 μm. (**b**) Quantification of RPA1 or RAD51 foci in MCF7 cells. A cell with > 5 distinct RPA1 or RAD51 foci in the nucleus was considered as positive. Error bars, SEM (n = 3). p values were determined by one-way ANOVA and Dunnett’s multiple comparisons test (**p < 0.01). (**c, d**) Representative confocal microscopy images of *KILR* and RPA1 in MCF7 (**c**) or MCF10A (**d**) cells after doxycycline induction of ectopic *KCTD1-5* or *KILR* expression and exposure to 6-Gy IR (n = 3). 6 h post-IR, cells were stained with Stellaris *KILR* RNA FISH probes (red) and immunostained with anti-RPA1 (green). The Tet-CON represents an empty vector control. Nuclei were stained with DAPI (blue). White arrows highlight *KILR*/RPA1 co-localization. Scale bar, 5 or 10 μm. (**e**) Representative confocal microscopy images of ɣH2AX in MCF7 cells after Cas13d-*KILR* knockdown with two independent crRNAs (crKILR1-2) and exposure to 6-Gy IR (n = 3). 6 h post-IR, cells were immunostained with anti-ɣH2AX (red). The crCON contains a non-targeting control. Nuclei were stained with DAPI (blue). Scale bar, 10 μm. Quantification of ɣH2AX foci in MCF7 cells. A cell with > 10 distinct ɣH2AX foci in the nucleus was considered as positive. Error bars, SEM (n = 3). p values were determined by one-way ANOVA and Dunnett’s multiple comparisons test (*p < 0.05, **p < 0.01).

## DISCUSSION

Although the number of lncRNAs has surpassed protein-coding genes, it is still unclear what proportion of lncRNAs are functional as opposed to transcriptional noise. High-throughput pooled CRISPR screens provide an unbiased method of identifying protein-coding and noncoding genes that function in different biological processes. CRISPR-Cas9 screens are commonly used to assess protein-coding genes for function, however they are often ineffective for lncRNAs as it is difficult to predict the impact of a Cas9-induced indel on lncRNA function. CRISPRi screens have successfully been used to identify functional lncRNAs^8,27,28^, but as they are often transcribed from enhancers, the observed phenotype can be a consequence of CRISPRi-induced enhancer suppression. Additional experiments are required to decipher if the phenotype is mediated by DNA or RNA.

Here, we describe the use of Cas13d-mediated RNA knockdown screens to identify breast cancer-associated lncRNAs that modulate proliferation in normal breast and breast cancer cells. Cas13d has previously been used to screen circular RNAs (circRNAs), with crRNAs designed to target their back-splicing junction allowing the discrimination of circRNAs from their host mRNA^29^. Several circRNAs were identified as important mediators of cell growth, including *circFAM120A* which was shown to promote cell proliferation by preventing the translation inhibitor IGF2BP2 from binding its host mRNA, *FAM120A* (and other family members^29^). Pooled Cas13d screens have also been used to optimize crRNA design. For example, fluorescent sorting for the cell surface markers CD46, CD55 and CD71 were used to screen for the best crRNA sequences for mRNA knockdown^30^. Using knowledge gained from these studies, Wessels *et al*^30^ developed a computational pipeline for crRNA design (*cas13design*), which we utilized in this study. Based on *cas13design* results, we selected the top ten non-overlapping crRNAs with the highest predicted quality for each lncRNA in the screen.

Off-targeting effects are one of the major limitations with CRISPR-based technologies and one that is often overlooked. To increase the likelihood of obtaining *bona fide* hits, we removed crRNAs with complementarity to genomic regions which were not the intended target. Optimally, this filtering step should be performed as part of the crRNA design prior to library synthesis. A limitation of using Cas13d in CRISPR screens is its reported collateral activity, where in addition to specifically cleaving the target RNA, it also promiscuously cleaves bystander RNAs^31,32^. To mitigate the consequences of this collateral activity, we individually validated the prioritized proliferation-related lncRNAs identified in our screens using multiple methods, prior to any follow-up characterization. Recently, a high-fidelity Cas13 enzyme was engineered (hfCas13d) which potentially minimizes collateral degradation of bystander RNA^33^. We anticipate that future CRISPR screens will benefit from the improved enzyme, representing an important advancement to the field.

The majority of GWAS variants are located in noncoding regions, frequently at lncRNA exons^7^, but there is limited functional evidence implicating lncRNAs in disease risk. In this study, we identified 43 lncRNAs (39 unannotated^7^ and four annotated) whose knockdown modulated breast cell proliferation, a fundamental trait of cancer cells. One the unannotated lncRNAs, *KILR* is transcribed from an intron of *KCTD1-5*, using an alternative promoter of *KCTD1*. As *KILR* is polyadenylated, it is likely to be a product of alternative polyadenylation rather than recursive mRNA splicing. Similarly, the start of *KILR* is ∼105 kb from the *KCTD1-5* TSS, indicating that its 5’ end is post-transcriptionally processed and stabilized. Our RNA folding predictions suggest that *KILR* possesses complex secondary structures at both terminal ends, which could explain how the transcript is protected from exonucleases. As *KILR* also has three predicted snoRNAs within its sequence, it is possibly a SPA-lncRNA (5’ small nucleolar RNA capped and 3′ polyadenylated), where the 5’ end is stabilized by a snoRNA structure rather than an m7G cap^34^.

The *KILR* breast cancer risk signal at 18q11 is colocalized with the genetic signal of the eQTL, suggesting that the risk variants can function by modulating *KILR* expression. In support of this, we show that the half-life of *KILR* in the presence of the risk alleles is significantly reduced as compared to that of *KILR* with the protective alleles. It is likely that one or more of the risk variants disrupts *KILR* secondary structure or affects the binding of a protein(s) responsible for maintaining *KILR* stability. This is the first time that GWAS variants have been shown to act by directly altering the RNA stability of a lncRNA transcript reducing its expression. Mechanistically, we showed that reduced *KILR* expression promoted breast cancer cell proliferation by increasing DNA replication fork speed. *KILR* binds to and sequesters RPA1, suggesting that its reduced expression would increase the available pool of RPA1. Previous studies have shown that the RPA complex participates in the initiation and elongation steps of DNA replication^35,36^ and that increased levels of RPA1 accelerate DNA replication and therefore promote cell proliferation^37^.

Overexpression of *KILR* mimics the reported effects of RPA1 knockdown on cancer cell growth^38^. In line with this, RPA1 deficiency has been shown to cause spontaneous DSBs and apoptosis^39^. Breast cancer cells partially depleted of RPA1 by siRNA treatment also become over-sensitive to DNA damage^40^. Indeed, relative expression of RPA is a predictor of response to chemotherapy in many cancers^41^. In breast cancer, RPA has also been linked with tumor aggressiveness and a decrease in overall survival^40^. Attempts to inhibit the RPA complex with synthetic molecules have resulted in cell death via apoptosis and has been established as a novel class of broad spectrum anticancer agents (RPAis^41^). The most promising first generation RPAi (TDRL-551) increases the efficacy of platinum-based chemotherapy in ovarian cancer^42^. Although this class of drugs was successful in preclinical studies, the RPAis explored so far presented chemical liabilities that could hinder their clinical use. RNA-focused therapy that interferes with cell proliferation and apoptosis has been cited as a promising avenue for cancer treatment^43^. We suggest that *KILR* could be used as an endogenous replacement of chemical RPAis or in combination therapy, if second generation synthetic RPAis prove to be safe.

We showed that overexpression of *KILR* in normal breast cells did not induce apoptosis. Normal cells are not subject to replication stress, which is often detected in highly-proliferative cancer cells and thus are less dependent on RPA availability. We observed a pool of free RPA1 in the normal breast cells which we hypothesized was sufficient to maintain DNA replication even after *KILR* overexpression. This idea is supported by the fact that after IR exposure, normal breast cells overexpressing *KILR* form RPA1 puncta independent of *KILR*, suggesting that free RPA1 molecules can aggregate (likely at DSBs) in response to DNA damage. In cancer cells, the combination of replication stress and defective DNA damage repair results in replication catastrophe and cell death.

RPA is critical to prevent this from happening as the exhaustion of free RPA1 leads to the accumulation of unprotected ssDNA and subsequent DSBs^44^. Further understanding the different outcomes of altering *KILR* expression in normal breast versus cancer cells will be important to determine the clinical relevance of *KILR*.

In summary, we have used high-throughput CRISPR-Cas13d screens to functionally evaluate lncRNAs at breast cancer risk regions identified by GWAS. This is the first time a large-scale CRISPR-Cas13 screen has been performed to efficiently target cancer-associated lncRNAs. For one lncRNA, *KILR* the presence of a GWAS variant resulted in its destabilization and reduced expression, which represents a new mechanism of action which may help explain a fraction of the GWAS. Mechanistically, reduced *KILR* expression promoted cell proliferation by increasing the pool of free RPA1 and speeding up DNA replication. *KILR* overexpression elicited apoptosis specifically in breast cancer cells, mediated by the binding and sequestration of RPA1 in nuclear foci. This likely depletes the cancer cells of available RPA culminating in impaired DNA replication and repair mechanisms. These latter results suggest that *KILR* could be explored as a novel RPA inhibitor, either replacing the synthetic drugs currently under development or working synergistically to potentialize their effect.

## METHODS

### Cell lines and culture

MCF7, MDAMB231, T47D, MCF10A, Hs578T and HEK293 cell lines were obtained from ATCC and grown according to their guidelines. MDAMB361 cells (ATCC) were grown in DMEM (Gibco Invitrogen) with 20% fetal bovine serum (FBS; Gibco Invitrogen) and 1% antibiotics (Gibco Invitrogen). B80T5 cells (a gift from Roger Reddel; CMRI, Australia) were grown in RPMI (Gibco Invitrogen) with 10% FBS and 1% antibiotics. K5+/K19+ cells^12^ were grown in 1:1 MEM α (Gibco Invitrogen) and Ham’s F-12 Nutrient Mix (Gibco Invitrogen) with 1% FBS, 10mM HEPES, 1 μg/ml bovine pancreatic insulin, 1 μg/ml hydrocortisone, 50 μg/ml epidermal growth factor (Sigma Aldrich), 10 mg/ml transferrin, 100 μM β-estradiol, 2 mM glutamine, 2.6 ng/ml sodium selenite, 1 ng/ml cholera toxin (Sigma Aldrich), 6.5 ng/ml triiodothyronine, 100 μM ethanolamine, 35μg/ml BPE, 10 μg/ml gentamicin, 10 μg/ml ascorbic acid, 15 μg/ml hygromycin B. All cell lines were tested for mycoplasma contamination and verified by short tandem repeats (STR) profiling.

### Plasmid constructs

To generate a Cas13d-NLS expression vector (pLXTRC311/NLS-EF1a-RxCas13d-2A-EGFP-blast; abbreviated to Cas13d), the Cas13d-NLS cassette was PCR-amplified from the pXR001_EF1a-CasRx-2A-EGFP (Addgene #109049) plasmid and cloned into pLX_TRC311-NLS-Cas13b-NES-P2A-Blast-eGFP. CRISPR-Cas13d crRNAs were cloned into BsmBI-digested pLentiRNAGuide_001 vector (Addgene #138150) and CRISPRa gRNAs into BsmBI-digested pXPR502 vector (Addgene #96923). For overexpression, full-length *KILR* was amplified from T47D cDNA using the KAPA HiFi PCR Kit (Kapa Biosystems) and *KCTD1-5* cDNA was synthesized by IDT. PCR products were cloned into the doxycycline-inducible plasmid pCW57-MCS1-2A-MCS2 (Addgene #71782), which was modified by adding bGHpolyA between the MluI and BamHI restriction sites. For pull-down assays, full-length *KILR* cDNA was cloned into pGEM-T (Promega). All constructs were confirmed by Sanger sequencing at the Australian Genome Research Facility (AGRF). The primers, crRNAs and gRNAs used in this study are provided in **Supplementary Table 4**.

### Generation of stable cell lines

Lentiviral plasmids were co-transfected with VSV-G envelope plasmid, pMD2.G (Addgene #12259) and packaging plasmid psPAX2 (Addgene #12260) into HEK293 using FuGENE 6 transfection reagent (Promega). Culture supernatant containing lentiviral particles was harvested after 24-48h incubation and passed through a 0.45 μm filter. Lentivirus was concentrated by centrifuging at 10,000 rpm at 4°C for 16-24h,resuspended in RPMI 1640 medium with 10% FBS, aliquoted and stored at –80°C. Breast cells were transduced at a high multiplicity of infection (MOI) with either Cas13d or CRISPRa (dCas9-VP64; Addgene #61425) lentivirus by spinoculation at 2,500 rpm for 1.5h at room temperature. To increase transduction efficiency, 5-8 μg/ml of polybrene (Sigma Aldrich) was supplemented in the media. Forty-eight hours post-transduction, cells were stabilized with 10-15 μg/ml blasticidin (Thermo Fisher Scientific) for two weeks and then maintained at 5-10 μg/ml blasticidin. Cas13d-expressing cells with high GFP were further purified by fluorescent activated cell sorting (FACS; FACSAria™ III Cell Sorter; BD Biosciences).

### CRISPR-Cas13 guide RNA (crRNA) library design

Cas13 crRNAs were designed using the basic algorithm in the *cas13design* tool^30^ (https://cas13design.nygenome.org) and further filtered to improve library quality. The following steps were followed: (1) Only the pool of high-quality guides (top quartile of quality scores) was considered for further analysis, unless step 4 is activated. (2) From the high-quality guides, we selected those with no overlap, to increase gene coverage. (3) According to the quality scores provided by *cas13design*, the top ten guides that meet the above criteria were selected per transcript. (4) For transcripts with less than five guides after filtering, we relaxed some of the criteria (e.g. allowing guides with quality scores in the third quartile or with 5-10 nucleotides overlap with each other, in this order). (5) We then re-run steps 2-3 for this subset and re-enter step 4 if necessary. (6) When all transcripts have 5-10 guides that pass the quality filtering described above, we stopped reiterating. (7) We removed redundancy in the library and added the required flanking sequences before sending the library to be synthesised (see below). (8) Blast alignments were used to remove guides with off-targets to either the reference human genome (hg38) or transcriptome (Gencode v.36). All crRNAs matching any region outside the target gene with up to two mismatches were considered as off-targets and removed from the *in silico* library.

### crRNA library generation

The oligonucleotides for the crRNA library were synthesized by Genscript. The sequences were collectively amplified with primers that generated 40 bp homologies with the pLentiRNAGuide_001 vector digested with BsmBI and XhoI. PCR was performed using Q5 High-Fidelity DNA Polymerase (New England Biolabs) for 20 cycles. The amplified crRNA library was then gel purified and assembled into BsmBI/Xhol-digested pLentiRNAGuide_001 using NEBuilder HiFi DNA Assembly master mix (NEB). The assembled plasmids were purified and concentrated by isopropanol precipitation. Three hundred nanograms of purified plasmids were electroporated into 25 µl of Endura electrocompetent cells (Lucigen) according to the manufacturer’s instructions. The electroporated cells were recovered in recovery medium (Lucigen) for 1 h and then plated on Terrific Broth (TB) agar plates with 100 μg/ml ampicillin at 37°C for 16 h. The resulting colonies were scraped and harvested in bulk at a coverage of more than 500 colonies per crRNA. The library plasmids were extracted using the NucleoBond Xtra Maxi EF Kit (Macherey-Nagel) to avoid endotoxin contamination. Library quality was assessed by next-generation sequencing.

### Pooled CRISPR-Cas13d proliferation screens

K5+/K19+, MCF7 and MDAMB231 cells stably expressing Cas13d were transduced with the crRNA library at an MOI of 0.3 to obtain 1000 cells/crRNA (three biological replicates per cell line). Twenty-four hours post-infection, cells were selected using 1-2 μg/ml puromycin and then maintained with 1-2 μg/ml puromycin and 10 μg/ml blasticidin throughout the screen to ensure crRNA and Cas13d expression. At 21 days post-infection, gDNA was extracted from the cells using the Quick-DNA Midiprep Plus Kit (Zymo Research), and one-step PCR was performed to amplify and add barcodes to the integrated crRNA sequences. PCR products were gel purified and sequenced by next-generation sequencing (20M reads/replicate). Quality control using FastQC v.0.11.8 (https://www.bioinformatics.babraham.ac.uk/projects/fastqc) was performed on the sequenced libraries and abundance estimation of all crRNAs using BBduk v.2019 (https://sourceforge.net/projects/bbmap) on Java v.1.8.192. Read counts were obtained for all crRNAs using MAGeCK v.0.5.9.4 run on Python v.3.6.1 and hits were called using MAGeCK test. A false discovery rate (FDR) threshold of 0.3 was applied to recover true hits in every cell line.

### Quantitative real-time PCR (qPCR)

Total RNA was extracted using TRIzol (Thermo Fisher Scientific). Complementary DNA (cDNA) was synthesized from RNA samples using SuperScript IV (Thermo Fisher Scientific). qPCR was performed using TaqMan assays (Thermo Fisher Scientific) or Syto9 incorporation into PCR-amplified products. Primers are listed in **Supplementary Table 4**.

### Cell proliferation assays

Cell proliferation was monitored using the IncuCyte live cell imaging system (Essen Bioscience). Cells were seeded at 2-3 x 10^4^ cells per well in 24-well plates and imaged using a 10× objective lens every 3 hours over 4-7 days. Imaging was performed in an incubator maintained at 37°C under a 5% CO_2_ atmosphere. Cell confluence in each well was measured using IncuCyte ZOOM 2016A software and the data analyzed using GraphPad Prism.

### Random amplification of cDNA ends (RACE)

5’ and 3’ RACE was performed using the GeneRacer kit (Thermo Fisher Scientific), following the manufacturer’s protocol. The purified PCR products were cloned into the pCR4-TOPO TA vector (Thermo Fisher Scientific) and identified by Sanger sequencing. Primers are listed in **Supplementary Table 4**.

### *KILR* secondary structure prediction and motif annotation

RNAfold, part of the Vienna package v.2.0^45^ was used for secondary structure predictions based on the *KILR* RNA transcript sequence (**Supplementary File 1**). The minimum free energy structure, based on the Turner model of 2004, was considered representative of *KILR*. The modeling temperature was defined as 37°C and isolated base pairs were avoided. The ALU elements that form the IRAlu structure of *KILR* were characterized based on Dfam v.3.6^46^ predictions. The machine learning algorithm implemented in snoReport v.2.0^47^ was used to identify snoRNA-like sequences in *KILR*. Other motifs such as the SIRLOIN nuclear localization sequence^19^ and BORG-like motifs were sourced from the literature.

### RNA stability assays

MDAMB361 cells were treated with 10 μg/ml actinomycin D (Sigma-Aldrich) to block transcription then harvested at 0, 3, 4, 8 and 12 hours post-treatment. qPCR was performed using a TaqMan™ Genotyping Assay (rs4555225 G/C; Thermo Fisher Scientific). A cyclin-dependent kinase inhibitor 2A (CDKN2A) TaqMan probe (Thermo Fisher Scientific) was used as an internal control. Linear regression analysis (GraphPad Prism) was used to estimate the decay rate of *KILR* with or without the risk alleles. The half-life was calculated by the equation t_1/2_ = ln(2)/k_decay_.

### Estrogen induction

Cells were treated with 10 nM fulvestrant (Sigma-Aldrich) for 48 h before the media was removed and replaced with media containing either 10 nM 17β-estradiol (Sigma-Aldrich) or DMSO (as vehicle control) for 24 h. Cells were harvested with TRIzol and assessed for induction of gene expression by qPCR.

### Cell fractionation

T47D cells were fractionated into subcellular compartments using the PARIS kit (Thermo Fisher Scientific). qPCR was performed to detect RNA in each fraction, with *RSP14* (Ribosomal protein S14), *U2snRNA* (U2 spliceosomal RNA) and *NEAT1* (nuclear enriched abundant transcript 1) serving as positive controls for RNA fractionated into the cytoplasmic, nuclear and chromatin compartments, respectively. Primers are listed in **Supplementary Table 4**.

### Apoptosis assays

For CRISPRa, cells were transduced with gRNAs and selected with 3 μg/ml puromycin and 10 μg/ml blasticidin for three days. Cells were then trypsinized, fixed and immunostained with the Alexa Fluor 488 Annexin V/Dead Cell Apoptosis Kit (Thermo Fisher Scientific), according to the manufacturer’s protocol, between 5-10 days post-transduction. For Tet-On overexpression, cells were transduced with lentivirus at a low MOI. Twenty-four hours post-transduction, the cells were treated with 1-3 μg/ml of puromycin for 4 days, then induced by 1 μg/ml doxycycline hyclate (Sigma-Aldrich) for 3 days, trypsinized, fixed and immunostained with the apoptosis kit according to the manufacturer’s protocol. The percentage of apoptotic cells was assessed by FACS_[HS1]_.

### RNA *in vitro* transcription and RNA-protein pull-down

The pGEMT-KILR construct was linearized with NotI then *in vitro* transcribed using the HiScribe T7 Quick High Yield RNA synthesis kit (NEB) according to the manufacturer’s instructions. LacZ RNA, produced from NotI-linearized pEF-ENTR-LacZ (Addgene #17430), was used as a negative control. RNA pull down was performed using the Pierce Magnetic RNA-Protein Pull-Down Kit (Thermo Fisher Scientific). Briefly, the *in vitro* transcribed RNAs were purified by TRIzol extraction, labeled with biotinylated cytidine bisphosphate, and incubated with cell lysates. After overnight incubation at 16°C, the RNA-protein complexes were captured with streptavidin beads and proteins were identified by mass spectrometry.

### Mass spectrometry

Samples underwent on-bead processing with 5 mM DTT at 60°C for 30 min then alkylated with 20 mM IAA for 10 min at room temperature in the dark. Proteins were digested with trypsin overnight at 37°C, then centrifuged at 20,000xg for 10 min to pellet the beads. The supernatants were acidified with trifluoroacetic acid, dried on a Speedvac, then reconstituted in 0.1% formic acid (FA) for LCMS analysis. Samples were loaded onto a Thermo Acclaim PepMap 100 trap column for 5 min at a flow rate of 10 μl/min with 95% Solvent A (0.1% FA in water) and separated on a Thermo PepMap100 analytical column equipped on a Thermo Ultimate 3000 LC interfaced with Thermo Exactive HF-X mass spectrometer. Peptides were resolved using a linear gradient of 5% solvent B (0.1% FA in 80% acetonitrile) to 40% solvent B over 48 min at a flow rate of 1.5 µl/min, followed by column washing and equilibration for a total run time of 65 min. Mass spectrometry data was acquired in positive ion mode. Precursor spectra (350-1400 m/z) were acquired on orbitrap at a resolution of 60,000. The AGC target was set to 3E6 with a maximum ion injection time of 30 ms. Top 20 precursors were selected for fragmentation in each cycle and fragment spectra were acquired in orbitrap at a resolution of 15,000 with stepped collision energies of 28, 30 and 32. The AGC target was 1E5, with a maximum ion injection time of 45 ms. The isolation window was set to 1.2 m/z. Dynamic exclusion was set to 30 s and precursors with charge states from 2-7 were selected for fragmentation. MS/MS data were searched against the reviewed Uniprot human database using Sequest HT on the Thermo Proteome Discoverer software (v.2.2). An FDR of 1% was used to filter peptide spectrum matches (PSMs). Carbamidomethylation of cysteines was set as a fixed modification, while oxidation of methionine, deamidation of glutamine and asparagine were set as dynamic modifications. Protein abundance was based on intensity of the parent ions and data were normalized based on total peptide amount. Five biological replicates were independently analysed for statistical significance, calculated using a *t*-test for summed abundance based ratios. Only proteins with at least five identified peptides, log2 [fold-change] (over LacZ) > 2.0 and p-value < 0.05 were considered. The resulting metrics were combined and the fold-change averaged across the replicates to obtain the final ranking of *KILR* protein partners.

### RNA-fluorescence *in situ* hybridization (FISH) and immunofluorescence (IF)

For RNA FISH, CRISPRa and Tet-On overexpressing cells grown on coverslips were fixed in 4% formaldehyde for 10 min followed by permeabilization in 70% ethanol overnight at 4°C. Cells were then stained for 16 h with 125 nM of a custom *KILR* Stellaris RNA-FISH probe set labelled with Quasar 570 fluorophore (LGC Biosearch Technologies) according to the manufacturer’s instructions. For IF, CRISPRa, CRISPR-Cas13 and Tet-On overexpressing cells were challenged with or without 6 Gy gamma irradiation followed by 6 h of incubation. The cells were then treated with CSK buffer (10 mM PIPES, 100 mM NaCl, 300 mM sucrose, 3 mM MgCl_2_, 1.4% Triton X-100) to remove the cytoplasm, followed by fixation in 4% formaldehyde for 10 min and permeabilization with 0.5% Triton X-100 for 15 min. The cells were incubated with antibodies against RPA1/RPA70 (Abcam, ab79398, 1:250), γH2AX (Abcam, ab2893, 1:1000) or RAD51 (GeneTex, GTX70230, 1:500). Coverslips were mounted onto slides using ProLong Glass antifade medium containing NucBlue nuclear counterstain (Thermo Fisher Scientific). Images were acquired with the DeltaVision Deconvolution microscope (GE Healthcare) using a 60× objective lens and analyzed with ImageJ software. A minimum of 100 cells per sample were analyzed.

### DNA fiber assays

CRISPR-Cas13d or Tet-On overexpressing cells were sequentially pulse-labeled with 50 µM 5-chloro-2′-deoxyuridine (CldU, Sigma-Aldrich) and 250 µM 5-iodo-2′-deoxyuridine (IdU, Sigma-Aldrich) for 25 min each, followed by treatment with or without 4 mM hydroxyurea (Sigma-Aldrich) for 4 h. Labeled cells were then washed and harvested in phosphate-buffered saline. Cell lysis, DNA spreading, denaturation and immunostaining were performed as described previously^48^. The slides were stained overnight at 4°C with anti-BrdU (Abcam, ab6326, 1:300) for CldU tracks and anti-BrdU (BD Biosciences, 347583, 1:50) to detect IdU tracks. After washing three times, the slides were stained for 1 h at 37°C with Alexa Fluor 488-labeled chicken anti-rat IgG (Invitrogen, A21470, 1:300) and Alexa Fluor 546-labeled goat anti-mouse IgG (Invitrogen, A11030, 1:300) secondary antibodies, respectively. Slides were visualized using the DeltaVision Deconvolution microscope with a 40× objective lens and analyzed with ImageJ software. A minimum of 150 fibers per sample were analyzed.

## AUTHOR CONTRIBUTIONS

L.W., X.L., S.N., H.S., S.J., K.M.H., S.K., F.C. and R.Z. performed the lab-based experiments. M.B. designed the Cas13d crRNA library and performed all the bioinformatic analyses. H.G.’s laboratory performed the mass spectrometry. J.R. assisted with the design of the study. J.D.F. and S.L.E. conceived and directed the study. L.W., M.B. S.L.E and J.D.F., wrote the manuscript with contributions from all authors.

## ACKNOWLEDGEMENTS

This work was supported by grants from Tour de Cure (RSP-225-1819), the National Health and Medical Research Council of Australia (1122022) and the National Breast Cancer Foundation of Australia (IIRS-23-060). L.W. was supported by a Queensland University of Technology (QUT) PhD scholarship. S.L.E. was supported by a NHMRC Senior Research Fellow (1135932). J.D.F. was supported by a philanthropic donation from Isabel and Roderic Allpass and an NHMRC Investigator Grant (2011196). The results published here are in part based upon the data generated by the TCGA Research Network. We thank Dr. Tetsuro Hirose for sharing his expertise on phase separation and lncRNAs. Funding for open access charge: QIMR Berghofer offers institutional grants.

